# Conformational dynamics of plasmepsin X during inhibitor binding

**DOI:** 10.64898/2026.01.06.697907

**Authors:** Wilson Karubiu, Richard Kullmann, Michael Krummhaar, Christian Roth, Thomas R. Weikl

**Author notes:** Centre for Targeted Protein Degradation, School of Life Sciences, University of Dundee, UK.

## Abstract

The aspartic protease plasmepsin X (PMX) of the parasite *Plasmodium* is a promising drug target for novel malaria therapies. Two potent inhibitors of PMX are WM382 and WM4, which both include a guanidinium group that is in contact with the two catalytic aspartates of PMX in the bound complexes. In structural representations of the inhibitors, the guanidinium group is typically depicted as uncharged. However, p*K*_*a*_ predictions with standard tools presented in this article indicate that the guanidinium groups of WM382 and WM4 are protonated and, thus, positively charged in the bound complexes. This positive charge is counterbalanced by a negatively charged catalytic aspartate D266 in PMX of *Plasmodium falciparum*, while the second catalytic aspartate D457 is uncharged. To investigate the interplay of the conformational dynamics of PMX and inhibitor (un)binding, we performed Hamiltonian replica exchange molecular dynamics (H-REMD) simulations starting from the predicted protonation state of the PMX-WM382 complex. On eight unbinding pathways enabled by weakened interactions of PMX and WM382 in the H-REMD simulations, we observed a dominant route of exit of the inhibitor from the binding pocket with a coupling to conformational changes in the “flap” of PMX, a *β*-hairpin located above the binding pocket. In the bound complex, the flap adopts a closed conformation in which it tightly interacts with and covers the inhibitor. On the dominant route observed in our simulations, unbinding involves an open conformation of the flap that allows the inhibitor to exit the binding pocket. After unbinding, the flap adopts an occluded conformation in which the binding site is blocked by a bulky aromatic sidechain.

## 1 Introduction

The ten plasmepsins I to X are aspartic proteases that are expressed at different stages of the lifecyle of the malaria-causing parasite plasmodium ^1,2^. In 2017, plasmepsin X was suggested as a promising drug target because of its essential role in the lifecyle ^3,4^. Two potent inhibitors of plasmepsin X are the small molecules WM4 and WM382^5^, which both contain a guanidinium group. In X-ray crystal structures of PMX in complex with WM382 and WM4, the guanidinium group of the inhibitors faces the two aspartates of plasmepsin X that constitute the catalytic dyad ^6^ (see Fig. 1). These aspartates are highly conserved in the plasmepsin family and act as a general acid and base in catalyzing hydrolysis of peptide bonds in substrates ^7,8^. A characteristic structural feature of plasmepsin binding sites is a *β*-hairpin, called “flap”, that covers substrates and inhibitors in the bound complexes. In plasmepsin II, the flap adopts an open conformation in X-ray crystal structures of the unbound state in which the binding site is accessible to ligands ^9^. In plasmepsin X, in contrast, the flap adopts an occluded conformation in crystal structures of the unbound state in which a bulky aromatic sidechain of the flap occupies and, thus, blocks the binding site ^6^.

**Fig. 1.**
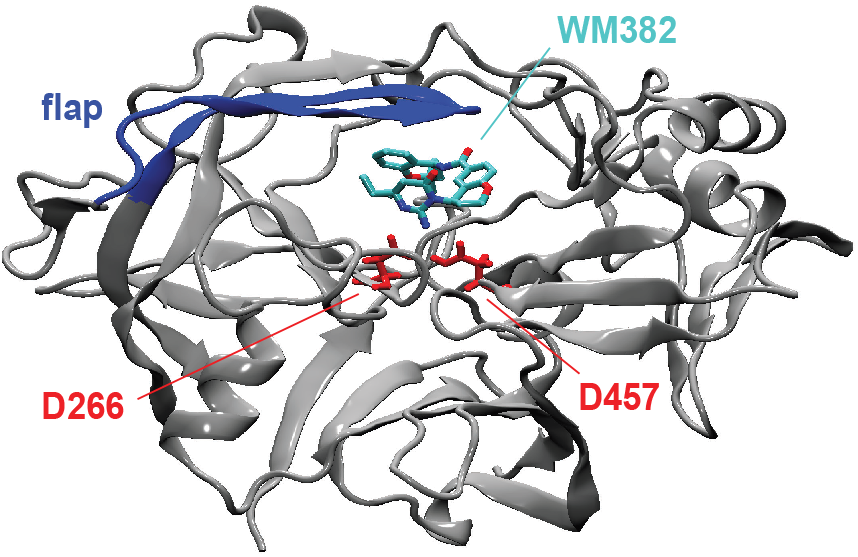
X-ray crystal structure of PMX from *Plasmodium falciparum* bound to WM382^6^. In this complex, WM382 is in contact with the two catalytic aspartates D266 and D457 and covered by the “flap”, a *β*-hairpin of PMX. The structure of PMX consists of two typologically similar N- and C-terminal subdomains, which form a crescent shape with a predominant *β*-sheet core. Each of the subdomains contributes one of the catalytic aspartates. A six-stranded interdomain *β*-sheet connects the two subdomains.

Here, we first determine the protonation states of the catalytic aspartates of plasmepsin X and of the guanidinium group of the inhibitors WM382 and WM4 in the bound complexes. These protonation states are not resolved in the X-ray crystal structures of the complexes due to the lack of hydrogen positions, but are required in molecular dynamics (MD) simulations, which we performed to investigate the coupling of the flap dynamics in plasmepsin X to inhibitor binding. For the unbound inhibitors, we obtain p*K*_*a*_ values of 7.7 and and 7.5 for the guanidinium group of WM382 and WM4 with MolGpka ^10^, a web server for small molecule p*K*_*a*_ predictions. To identify the shift of these p*K*_*a*_ values due to the chemical environment of the guaninidium groups in the complexes of WM382 and WM4 with plasmepsin X, we used PROPKA 3^11^, a widely used standard method for predicting p*K*_*a*_ shifts of the titratable groups of both proteins and bound ligands based on X-ray crystal structures. For WM382, PROPKA 3 predicts an upward shift of 3.7 of the p*K*_*a*_ value for the guanidinium group in the bound complex of WM382 and plasmepsin X. The p*K*_*a*_ of the WM382 guanidinium group thus shifts from 7.7 to about 11.4 during binding to plasmepsin X, which indicates that the guanidinium group is protonated. The positive charge of the protonated guanidinium group is counterbalanced by a negative charge of the catalytic aspartate D266 predicted by PROPKA 3, while the second catalytic aspartate D457 is predicted to be uncharged in the physiologically relevant pH range. The same protonation states are predicted for WM4 in complex with plasmepsin X.

To investigate the unbinding pathway of the inhibitor WM382 from plasmepsin X, we performed Hamiltonian replica-exchange MD (H-REMD) simulations. In these simulations, we observed inhibitor unbinding on sub-microsecond timescales accessible in atomistic simulations due to the exchanges to Hamiltonian replica energy levels on which the interactions between protein and inhibitor are strongly weakened, rather than on realistic timescales of seconds or minutes to be expected for inhibitors with a picomolar inhibition constant as WM382 (*K*_*i*_ = 38 *±* 8 pM) ^5^. Along the unbinding pathways in the H-REMD simulations, the flap of plasmepsin X first opens, which allows WM382 to leave the binding pocket. After unbinding, the flap adopts an occluded conformation in the simulations in which it blocks the binding site, in agreement with the crystal structure of unbound plasmepsin X ^6^. The H-REMD unbinding pathways are – strictly speaking – “unphysical” or “al-chemical”, because they involve unphysical, weakened interactions of the binding partners on Hamiltonian replicas visited along the pathways. However, we propose that the steric requirements suggested for inhibitor unbinding, with the flap opening prior to unbinding, also hold in the physical, realistic system.

## Methods

### System setup and relaxation

Because the N- and C-terminus of PMX are not resolved in the X-ray crystal structure of the PMX-WM382 complex (PDB ID: 7tbc), we capped the first and last PMX residue of the structure with Ace and Nme residues, respectively, and assigned the protonation states of the titratable protein residues at the physiological relevant pH of 4.5 with PROPKA 3^11^. We used the ff14SB force field ^12^ for the protein and parametrized WM382 with the General Amber Force Field 2 (GAFF2) and the AM1-BCC method for determining partial charges in the Antechamber package of the Amber20 software suite ^13^. We solvated the protein-inhibitor complex in TIP3P water in a truncated octahedron with the LEaP module of the Amber20 software suite for a 10 Å standard minimum distance between atoms of the protein complex and the edges of the box ^13,14^, adding 6 Na^+^ ions to attain charge neutrality of the simulation system. To mimic the physiological salt concentration of 150 mM, we further added appropriate numbers of Na^+^ and Cl^−^ ions for our simulation box size.

To relax the simulation system, we started with a standard two-step energy minimization approach. In the first minimization step, only water and ions were minimized in 5000 steepest descent steps followed by further 5000 conjugate gradient steps, while the protein-ligand complex was restrained with a harmonic force constant of 100 kcal mol^−1^ Å^−2^ on non-hydrogen atoms. In the second minimization step, all atom positions were minimized in 5000 steepest descent steps followed by further 5000 conjugate gradient steps. Next, we gradually heated the simulation system from 0 to 300 K in a 500 ps simulation in the NVT ensemble using a Langevin thermostat and harmonic restraints with a force constant of 100 kcal mol^−1^ Å^−2^ on the non-hydrogen atom positions of the protein-inhibitor complex. We further relaxed the system in a 4 ns simulation in the NPT ensemble at 300 K without any positional restraints.

### H-REMD simulations

We started the H-REMD simulation from initial conformations generated in five standard-MD trajectories of the complex after a simulation length of 1 µs. The H-REMD simulation with 20 replicas had a length of 1 µs and was performed on 20 GPUs in parallel with the pmemd.cuda package of the AMBER20 software suite ^13^. We used the repulsive-scaling H-REMD method introduced by Siebenmorgen, Engelhard, and Zacharias ^15^ in which the effective van der Waals radii and the van der Waals attraction in the Lennard-Jones interaction potential *V*_*i, j*_ of atoms *i* and *j* are modified by the parameters *d* and *e*:

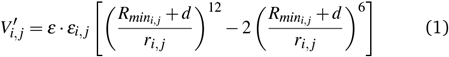

In our simulation, we employed the parameter values listed in Table 1, with additional cubic corrections of *ε*_*i, j*_ to compensate for effective increases of van der Waals interactions with increasing van der Waals radii ^15^. An isotropic pressure of 1 bar was maintained using a Berendsen barostat with a pressure relaxation time of 2 ps, the temperature was kept at 300K using a Langevin thermostat with a collision frequency of 1 ps^−1^, bonds containing hydrogen atoms were constrained using the SHAKE algorithm, a cutoff length of 10 Å was used for non-bonded interactions, and long-range electrostatic interactions were calculated with the Particle Mesh Ewald method. In addition, we used hydrogen mass repartitioning ^16^ to increase the simulation timestep to 4 fs. Exchanges between replicas were attempted at simulation intervals of 0.5 ps.

**Table 1.** Parameters *d* and *ε* of the rescaled Lennard-Jones potential (1) for the 20 replicas of our H-REMD simulations.

|  |  |  |  |  |  |  |  |  |  |  |
| --- | --- | --- | --- | --- | --- | --- | --- | --- | --- | --- |
| replica | 1 | 2 | 3 | 4 | 5 | 6 | 7 | 8 | 9 | 10 |
| $d$ | 0.0 | 0.03 | 0.06 | 0.09 | 0.12 | 0.15 | 0.18 | 0.21 | 0.24 | 0.28 |
| $\epsilon$ | 1 | 0.99 | 0.98 | 0.97 | 0.96 | 0.95 | 0.94 | 0.93 | 0.92 | 0.9 |
| replica | 11 | 12 | 13 | 14 | 15 | 16 | 17 | 18 | 19 | 20 |
| $d$ | 0.32 | 0.36 | 0.4 | 0.44 | 0.48 | 0.52 | 0.56 | 0.6 | 0.65 | 0.7 |
| $\epsilon$ | 0.88 | 0.86 | 0.84 | 0.82 | 0.8 | 0.78 | 0.76 | 0.74 | 0.72 | 0.71 |

## Results

### Protonation state prediction of the PMX-inhibitor complexes

In empirical p*K*_*a*_ predictions of titratable groups of proteins, a general strategy is to predict the shift of the p*K*_*a*_ values of these groups due to the chemical environment in the protein or protein complex, relative to reference p*K*_*a*_ values of the isolated groups in solvent. PROPKA 3 is one of the most widely used tools to predict the p*K*_*a*_ shifts of the titratable groups of both proteins and bound ligands based on X-ray crystal structures or other experimental structures of protein complexes that lack hydrogen positions ^11^. A complication in applying PROPKA 3 to the X-ray crystal structures of plasmepsin X in complex with WM382 and WM4 is that PROPKA 3 only recognizes guanidinium groups as terminal groups, assigning a reference p*K*_*a*_ value of 11.5 to such terminal guanidinium groups. In WM382 and WM4, however, the guanidinium groups are part of ring structures, and not terminal.

To assess the effect of the ring structure in WM382 and WM4 on the p*K*_*a*_ values of the guanidinium group, we use MolGpKa, a method for predicting p*K*_*a*_ values of small molecules ^10^. MolGpKa predicts a p*K*_*a*_ value of 12.7 for the guanidinium group as single group (see Fig. 2), which is somewhat larger than the reference value of 11.5 assigned by PROPKA 3. For a guanidinium group that is covalently bound to a carbonyl group as in the ring structures of WM382 and WM4, however, MolGpKa predicts a strong reduction of the p*K*_*a*_ value to 6.8. For the complete inhibitor WM382, MolGpKa predicts a p*K*_*a*_ value of the guanidinium group of 7.7 (see Fig. 2). The reduced p*K*_*a*_ value of the guanidinium group in WM382, compared to guanidinium as single group, thus appears to result mainly from the carbonyl group that is covalently bound to the guanidinium group. For WM4, MolGpKa predicts a p*K*_*a*_ value of the guanidinium group of 7.5, which is close to the p*K*_*a*_ value in WM382 because of the structural similarity (see Fig. 2).

**Fig. 2.**
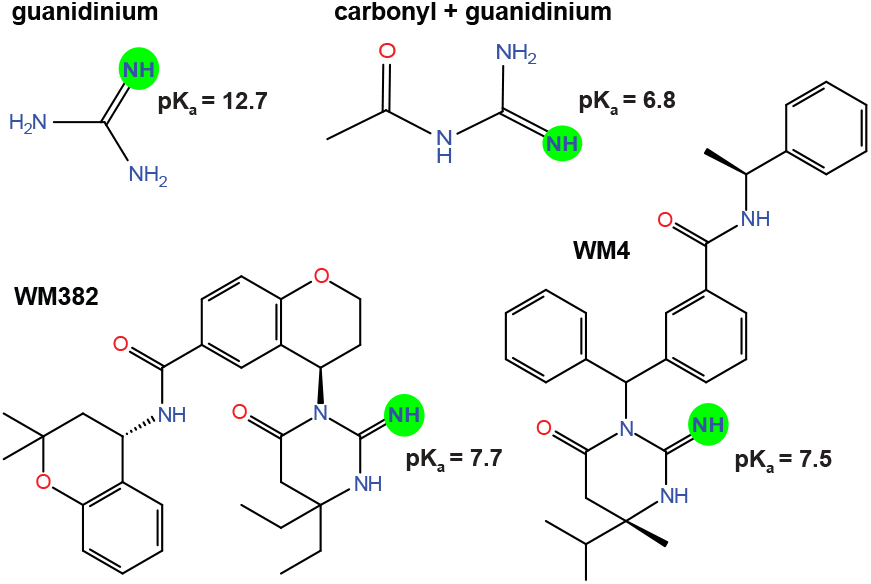
MolGpKa prediction of the p*K*_*a*_ of the isolated guanidinium group, of the guanidinium group bound to a carbonyl group, and of the guanidinium groups of the inhibitors WM382 and WM4.

Because PROPKA 3 does not recognize the guanidinium group in the ring structures of the inhibitors WM382 and WM4, we performed PROPKA 3 calculations with structural segments of the inhibitors. When we retain only the non-hydrogen atoms of the covalently bound guanidinium and carbonyl groups of the inhibitor WM382 in the X-ray crystal structure of plasmepsin X in complex with WM382, PROPKA 3 predicts a p*K*_*a*_ shift of +3.7. The same p*K*_*a*_ shift is predicted by PROPKA 3 when larger fragments of WM382 are retained in the crystal structure of the complex. Taking the MolGpKa-predicted guanidinium p*K*_*a*_ of 7.7 for the unbound inhibitor WM382 (see Fig. 2) as correct reference value, we obtain p*K*_*a*_ ≈ 7.7 + 3.7 ≈ 11.4 for guanidinium of WM382 in complex with plasmepsin X, which indicates that the guanidinium group is protonated. The positive charge of the protonated guanidinium group is counterbalanced by a negative charge of the catalytic aspartate D266 predicted by PROPKA 3, while the second catalytic aspartate D457 is predicted to be uncharged in the physiologically relevant pH range. The same protonation states are predicted for WM4 in complex with plasmepsin X.

### Unbinding pathways of WM382 in Hamiltonian replica exchange MD (H-REMD) simulations

To investigate the unbinding pathways of WM382 from PMX, we performed a H-REMD simulation starting from the relaxed complex in the protonation state predicted in the previous section (see Methods for details). The H-REMD simulation had a length of 1 µs and included 20 Hamiltonian replicas of the system, in which the interactions of plasmepsin X and WM382 are gradually weakened. Exchanges between neighbouring replicas were attempted every 0.5 ps and accepted or rejected with a standard exchange criterion that ensures detailed balance. From this H-REMD simulation, we constructed the 20 coordinate-continuous trajectories that are obtained by following the 20 initial conformations along its exchanges between the Hamiltonian replicas. WM382 unbound on 8 of these 20 coordinate-continuous trajectories. Figure 4 illustrates the root-mean-square-deviation of WM382 (RMSD_WM382_) along the 8 unbinding trajectories relative to the WM382 conformation in the crystal structure of the complex, after alignment of PMX. For RMSD values smaller than about 2 to 3 Å, WM382 adopts native-like bound conformations. For RMSD values smaller than about 15 to 20 Å, WM382 is still in contact with PMX in intermediate bound conformations. For RMSD values larger than 20 Å, WM382 is unbound. On 3 of the 8 coordinate-continuous unbinding trajectories (numbered 1 to 3 in Figure 4), WM382 reaches the unbound state from the initial native-like bound state within 15 to 30 ns. On trajectories 4 and 5 in Figure 4, WM382 unbinding occurs at about 80 and 110 ns, respectively. On the remaining trajectories 6 to 8, unbinding takes place between 200 ns and 400 ns. On several of the unbinding trajectories, WM382 stays in binding intermediate states with RMSD values smaller than 15 to 20 Å for some time prior to unbinding, e.g. on trajectories 5, 6, and 8.

**Fig. 3.**
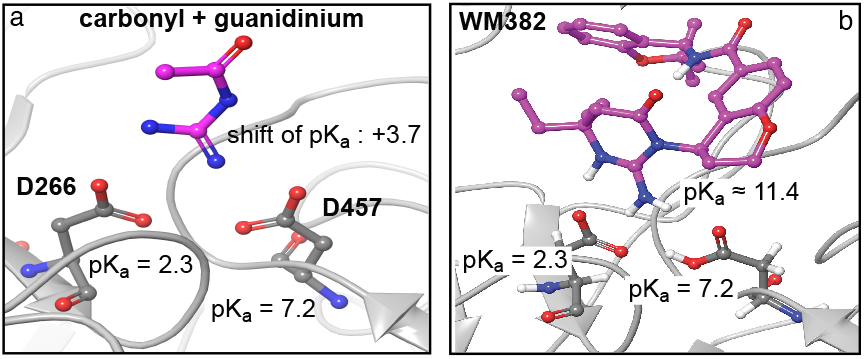
(a) PROPKA 3^11^ predicts a p*K*_*a*_ shift of +3.7 for the covalently bound guanidinium and carbonyl groups of the inhibitor WM382 in the Xray crystal structure of plasmepsin X in complex with WM382. (b) With the p*K*_*a*_ value of 7.7 predicted by MoGpKa ^10^ for the guanidinium group of the unbound inhibitor WM382, we obtain p*K*_*a*_ ≈7.7 + ≈3.7 11.4 for guanidinium of WM382 in complex with plasmepsin X. Hydrogens of nitrogen atoms in WM382 and at the sidechains of the catalytic aspartates D266 and D457 are depicted for the physiologically relevant pH range.

**Fig. 4.**
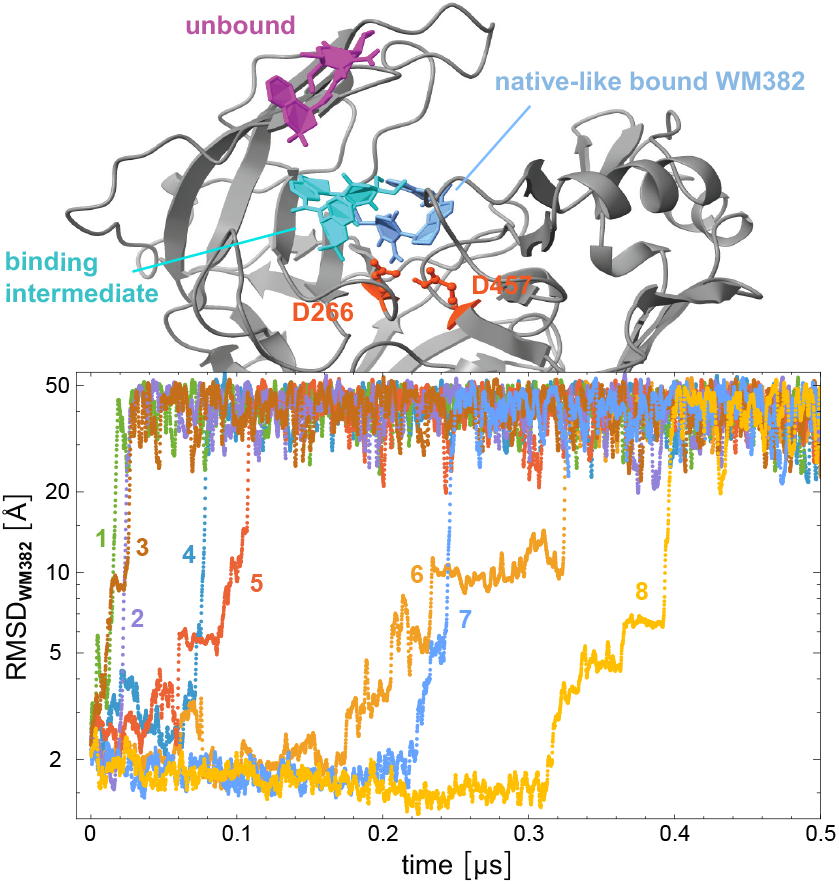
Root mean square deviation of WM382 (RMSD_WM382_) relative to the WM382 non-hydrogen-atom coordinates in the X-ray crystal structure of the complex with plasmepsin X along the 20 coordinate-continuous trajectories of the H-REMD simulation with 20 replicas. On 8 of the 20 trajectories, the ligand unbinds and attains large RMSD values relative to the native binding position. All observed unbinding events occur within the first shown 0.5 *µ*s of the H-REMD run with a total length of 1 *µ*s. Here, the RMSD_WM382_ values are smoothened by averaging over 11 frames at intervals of 0.1 ns. Representative snapshots of WM382 in the native-like bound state, in an intermediate state, and in the unbound state are shown in blue, cyan and magenta, respectively.

Along the 8 unbinding trajectories of our H-REMD simulation, WM382 unbinding occurs after an opening of the *β*-hairpin flap that covers the inhibitor in the native-like bound state. In Figure 5, the distance between C_*α*_ backbone atoms of the catalytic aspartate D266 and of phenylalanine F311 located at the tip of flap is plotted versus the RMSD value of WM382 along the 8 unbinding trajectories. At the smallest RMSD values of WM382, the C_*α*_ distance of D266 and F311 adopts values around 14 Å as in the crystal structure of the PMX-WM382 complex, in which this distance is 13.8 Å. In binding intermediate states with RMSD values of WM382 between about 5 and 20 Å, the C_*α*_ distance of D266 and F311 attains values up to 20 Å, on unbinding trajectory 6 even values up to 22 Å, that are clearly larger than the values around 14 Å in the native-like bound state and, thus, indicate an opening movement of the flap. After unbinding, i.e. for RMSD values of WM382 larger than 20 Å in Figure 5, the C_*α*_ distance of D266 and F311 predominantly adopts values smaller than 14 Å, which indicate a slight movement of F311 into the binding pocket, thus occluding the pocket. This occlusion of the binding pocket observed in our simulations after WM382 unbinding is in agreement with the crystal structure of unbound WM382, in which the C_*α*_ distance of D266 and F311 is 12.9 Å ^6^, and is accompanied by a reorientation of the terminal benzol ring of the F311 side chain. In Figure 6, the F311 ring orientation along our unbinding trajectories is quantified as the angle between the normal vector of the ring perpendicular to the ring plane and an axis that runs through the C_*α*_ atoms of D266 and F311. In bound states of WM382 with RMSD values smaller than about 15 Å, the F311 ring orientation angle attains values predominantly around 20°, which indicate that the perpendicular direction of the ring tends to be aligned to the axis connecting the C_*α*_ atoms of D266 and F311. In the unbound state, in contrast, the F311 ring orientation is distributed over the full range of possible angles between 0 and 90°, which indicates a dangling motion of the F311 ring that contributes to a steric blocking of the binding pocket.

**Fig. 5.**
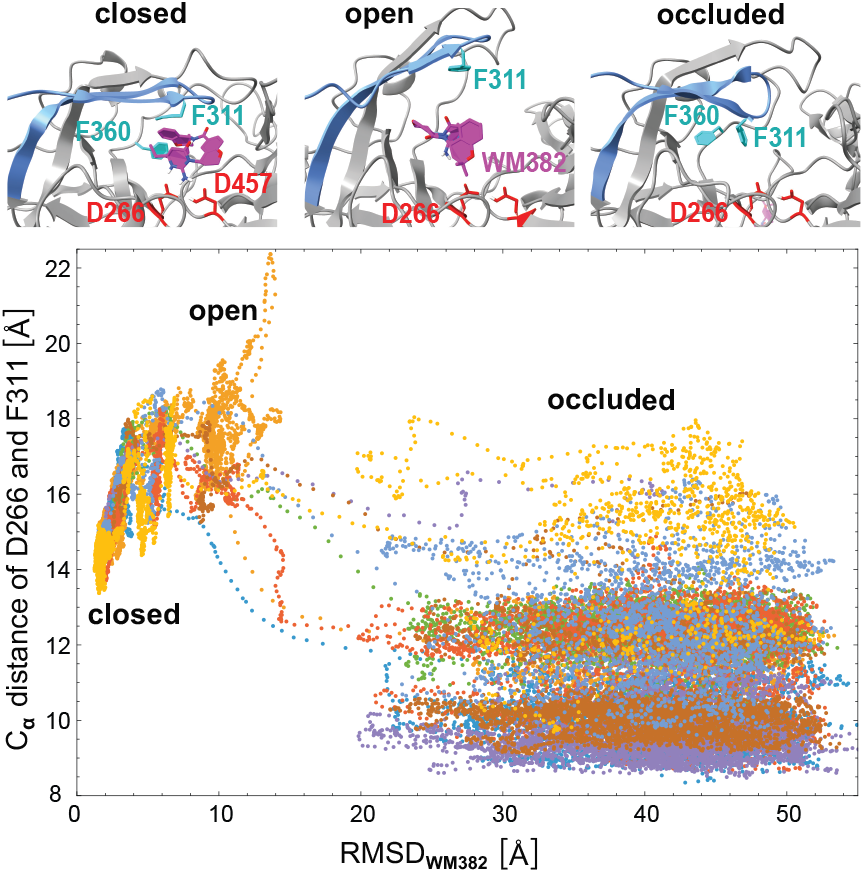
C_*α*_ distance of the catalytic aspartate D266 and phenylalanine F311 at the tip of flap versus the RMSD value of WM382 relative to the bound complex along the 8 unbinding trajectories of Figure 4. The C_*α*_ distances and RMSD values are averages over 11 simulation frames at intervals of 0.1 ns. In the exemplary conformations shown at the top, the C_*α*_ distance of D266 and F311 is 15.1 Å (closed), 20.5 Å (open), and 11.6 Å (occluded).

**Fig. 6.**
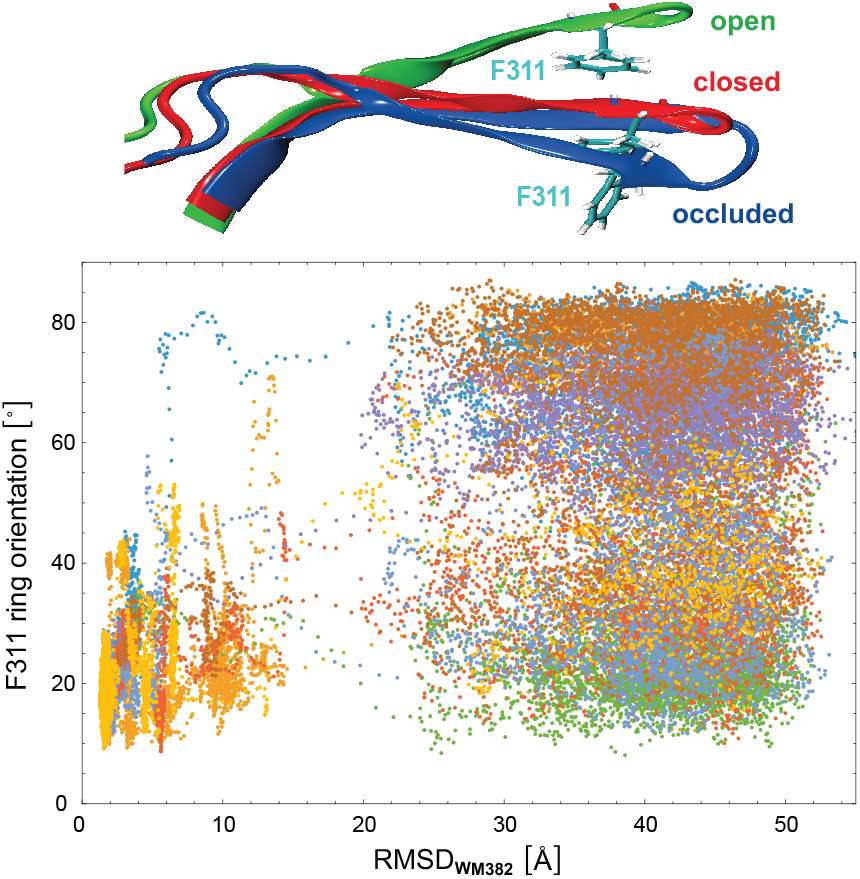
F311 ring orientation angle versus the RMSD value of WM382 relative to the bound complex along the 8 unbinding trajectories of Figure 4. Here, the F311 ring orientation angle is defined as the angle between an axis that is perpendicular to the plane of the terminal benzol ring of phenylalanice F311 and (b) and axis connecting the C_*α*_ atoms of D266 and F311. The F311 ring orientation angles and RMSD values are averages over 11 simulation frames at intervals of 0.1 ns.

## Discussion and conclusions

The unbinding pathways observed in our H-REMD simulations can be seen as “unphysical” or “alchemical” because they involve unphysical, weakened interactions of the binding partners on Hamiltonian replicas visited along the coordinated-continuous trajectories. However, it is important to note that the weakening of the interactions between the binding partners rather introduces a bias towards direct unbinding from closed conformations in situations where such direct unbinding is available as alternative route to unbinding after opening ^17^. Because we do not observe such direct unbinding from the closed-flap conformation along our eight unbinding trajectories for weakened interactions of the binding partners, we conclude that such direct unbinding also does not occur for the original physical interactions of the binding partners. In other words, the reasons for the flap opening prior to inhibitor unbinding observed along our unbinding trajectories appear to be steric, rather than energetic: The exit of the inhibitor appears to be sterically blocked by the closed flap, and is only possible if the flap opens, even at the strongly weakened interactions of our H-REMD scheme at which the inhibitor has a strong tendency to unbind.

Because thermodynamic principles imply that the binding pathway of PMX and WM382 is the reverse of the unbinding pathway, our simulation results indicate that WM382 can only bind to PMX conformations with an open flap. Both the closed and the occluded conformation of the flap appear to sterically block inhibitor access to the binding site. The conformational flap dynamics of PMX thus appears to affect also the binding rates, because binding in the pre-dominantly populated occluded conformation of the flap is not possible. Binding requires a open conformation of the flap, which implies that the binding rates are proportional to the relative population of the minor, open conformation of unbound PMX. For plasmepsin II, in contrast, the flap adopts an open conformation in a crystal structure of the unbound protein ^9^, which is also supported by MD simulations in which the flap is observed to remain open ^18,19^.

## Acknowledgment

WK thanks the DAAD and all authors thank the Max Planck Society for generous financial support. WK and RK thank Till Siebenmorgen for providing a script to carry out the rescaling of the Lennard-Jones potential (1).

